# Urine as a high-quality source of host genomic DNA from wild populations

**DOI:** 10.1101/2020.02.18.955377

**Authors:** Andrew T. Ozga, Timothy H. Webster, Ian C. Gilby, Melissa A. Wilson, Rebecca S. Nockerts, Michael L. Wilson, Anne E. Pusey, Yingying Li, Beatrice H. Hahn, Anne C. Stone

## Abstract

The ability to generate genomic data from wild animal populations has the potential to give unprecedented insight into the population history and dynamics of species in their natural habitats. However, in the case of many species, it is impossible legally, ethically, or logistically to obtain tissues samples of high-quality necessary for genomic analyses. In this study we evaluate the success of multiple sources of genetic material (feces, urine, dentin, and dental calculus) and several capture methods (shotgun, whole-genome, exome) in generating genome-scale data in wild eastern chimpanzees (*Pan troglodytes schweinfurthii*) from Gombe National Park, Tanzania. We found that urine harbors significantly more host DNA than other sources, leading to broader and deeper coverage across the genome. Urine also exhibited a lower rate of allelic dropout. We found exome sequencing to be far more successful than both shotgun sequencing and whole-genome capture at generating usable data from low-quality samples such as feces and dental calculus. These results highlight urine as a promising and untapped source of DNA that can be noninvasively collected from wild populations of many species.

## Introduction

The development of methods to generate genetic data from noninvasively collected samples revolutionized the study of wild animal populations, allowing for DNA research without the capture or even observation of species of interest (Kohn & Wayne, 1997; Waits & Paetkau, 2005). While studies of individual DNA markers improved our understanding of behavior, ecology, and evolution, recent advances in massively parallel sequencing strategies make it possible to incorporate information from across the entire genome, giving unprecedented insight into the evolution and population history of non-model species (Ellegren, 2014). However, for many species, it is impossible legally, ethically, or logistically to obtain high-quality tissue samples required for large-scale genomic analyses. It is therefore critically important to develop and evaluate methods for sampling and capturing genome-scale data from noninvasive and alternative sources.

While a variety of noninvasively collected biological materials have been used in DNA analyses, feces have been the primary target of recent attempts to generate genomic data. Rich in gut epithelial cells and often the most abundant, easiest to collect source of DNA in the environment, feces have long played a role in noninvasive genetic analyses (Constable, Ashley, Goodall, & Pusey, 2001; Hoss, Kohn, Paabo, Knauer, & Schroder, 1992; Kohn & Wayne, 1997). However, the retrieval of DNA from feces presents a number of difficulties. Challenges, including low DNA yields, DNA fragmentation and degradation (Deagle, Eveson, & Jarman, 2006) and the presence of PCR inhibitors, can lead to genotyping errors (Taberlet, Waits, & Luikart, 1999). Moreover, DNA recovered from fecal material is dominated by microbes (>95% exogenous DNA), which further complicates genotyping (Perry, Marioni, Melsted, & Gilad, 2010). For genetic analyses involving small number of markers, these challenges are well understood and can be overcome. However, these problems are exacerbated in massively parallel sequencing, which typically requires higher quantities and qualities of input DNA, and generates almost entirely microbial data due to the very low levels of host DNA in samples.

The main strategy that has been employed to combat these problems is enrichment of host DNA. In this vein, there have been three major methodological developments. Perry and colleagues (2010) first enriched DNA from feces on a genomic scale by using custom chimpanzee baits designed to capture approximately 1.5 Mb of sequence across six western chimpanzees (*Pan troglodytes verus*). While successful, this method required a reference genome to design baits and was cost prohibitive for producing genome-scale datasets. To address these challenges, Snyder-Mackler and colleagues developed a protocol to create RNA baits from high-quality host DNA and improve post-capture enrichment (2016). However, for this method, bait requirements—notably high-quality host DNA—and low sequencing coverage of host DNA remain barriers for some study systems and questions. Recently, Chiou and Bergey introduced a method that exploits differences in CpG-methylation densities between vertebrate and bacterial genomes to capture host DNA, alleviating the need for high-quality host material or reference genome to design baits (2018). However, CpG content varies substantially across the genomes of primates and other mammals (Han, Su, Li, & Zhao, 2008), thus targeting these regions specifically may bias the regions captured.

Despite these improvements to both capture and enrichment, DNA capture from feces is still far less efficient than from high-quality tissues. This leads to a tradeoff: attempting to capture large genomic regions leads to very low sequence coverage; however, targeting a subset of the genome can lead to biases. A compromise would be to target a small subset of the genome that is biologically important. One potential option is exome sequencing, a capture-based method that targets the entire coding region of the genome, comprising approximately 1.5% of the total length of the genome. Coding regions are among the best understood in the genome and are of great evolutionary and conservation interest (Bataillon et al., 2015; George et al., 2011; Hvilsom et al., 2012). Because exome sequencing is so widely used in human genomics, many commercial kits are available and much cheaper than custom alternatives. Human exome baits have been successfully used in a number of nonhuman primate studies (George et al., 2011; Jin et al., 2012; Vallender, 2011) and have been shown to work in primate species as distantly related from humans as Strepsirrhines (Webster, Guevara, Lawler, & Bradley, 2018). Moreover, recent work has shown that exome capture successfully enriches host DNA in chimpanzee fecal samples (Hernandez-Rodriguez et al., 2018; White et al., 2019). However, some of this work involves first screening for endogenous content using quantitative PCR (qPCR), which although successful, can be a limiting factor for smaller labs at the scales for population genomics. For example, after screening 1,780 fecal samples, White and colleagues estimated 101 samples contained enough endogenous DNA for sequencing (>1%) (White et al., 2019).

In addition to methodological development, turning to other sources of biological material might improve sequencing success in wild populations. Efforts up to this point have focused almost exclusively on feces, and many other noninvasive alternatives remain to be explored. Urine, in particular, is abundant for many large-bodied species, and has been used, albeit infrequently, as a source of DNA collected noninvasively from the environment (Hedmark et al., 2004; Sastre et al., 2009; Valiere & Taberlet, 2000). Although difficult to obtain in certain field conditions, urine contains far fewer microbes than feces, does not contain traces of dietary DNA, and lacks many inhibiting compounds commonly found in feces that impact PCR success (Hausknecht, Gula, Pirga, & Kuehn, 2007; Inoue, Inoue-Murayama, Takenaka, & Nishida, 2007; Thomas-White, Brady, Wolfe, & Mueller, 2016). Another source of interest is skeletal material, which is often found at field sites and in museum collections. Dentin and dental calculus, in particular, are both capable of yielding host nuclear DNA (Ziesemer et al., 2018). Combining data from historic populations with those from contemporary populations has the potential to provide genomic insight into wild populations on a scale not yet fully realized.

In this study, we evaluate the success of several sources of host DNA and capture methods in generating genome-scale data in a population of wild, endangered animals. Specifically, we extracted and captured endogenous DNA from feces, dental calculus, dentin, and urine recovered from wild chimpanzees (*P. t. schweinfurthii*) from Gombe National Park, Tanzania. From these data we compared the success of whole-genome capture and targeted exome capture. We demonstrate that urine harbors the highest concentration of endogenous DNA of the materials sampled in this study. For other sources, whole-genome sequencing appears possible, but not cost-effective. Employing a targeted approach, such as exome capture, reduces the amount of sequence obtained in the genome, but it may result in increased sequencing efficiency. Finally, we show that genotypes generated from fecal and urine samples exhibit high levels of concordance and argue that genotypes from urine are less subject to contamination. Together, our results demonstrate that, while further methodological advances might improve host DNA extraction in feces, dentin, and dental calculus, urine is a promising source of noninvasive DNA from which genome-scale data can be easily generated. We anticipate the ability to generate genomic data from urine to be broadly useful across study systems, including many protected species.

## Materials and Methods

### Sample Collection and Extraction

We collected fecal samples in RNAlater from four wild chimpanzees as described (Stone et al., 2010) (7069, 7150, 7365, and 7507) from Gombe National Park between August 2011 and January 2014 and shipped them to the University of Pennsylvania for storage at −80°C. Using a sterile cut pipette tip, we removed roughly 200 μL of the fecal slurry and extracted DNA using QIAamp DNA Stool Minikit (Qiagen) according to manufacturer’s protocol. To obtain enough DNA, we repeated this process 8-12 times for each sample, then pooled and desiccated each sample down to 50-100 μL. We combined a total of 2 μg of DNA and molecular grade H_2_O into a 50 μl tube and then sheared DNA using a Covaris Sonicator for 4min at 150 bp according to manufacturer specifications.

We retrieved dental calculus from two skeletons (individuals 7057 and 7433; less than ten mg per sample) and dentin from one skeleton (individual 7057; less than 50 mg per sample) at the University of Minnesota using a sterile dental scaler. We decontaminated calculus using exposure to UV irradiation for five min. This was followed by an initial 0.5M EDTA (Ambion) wash in a 2.0 mL tube for 15 min. We subjected samples to a two day 0.5 EDTA and proteinase K (10 mg/mL; Qiagen) digestion, at which point we combined the resulting solution with 12 mL of PB buffer and followed standard MinElute PCR Purification Kit (Qiagen) protocol. Our dentin protocol followed previously published methods (Nieves-Colón et al., 2018). We did not shear dental calculus and dentin samples prior to shotgun library builds.

We collected urine from seven wild Gombe chimpanzees—three with matched fecal samples (7150, 7365, and 7507) and four others (7072, 7323, 7535, and 7650)—in the early morning using fresh plastic bags attached to sticks suspended below chimpanzee nests. We immediately transferred between 10 mL and 30 mL of urine to a 50 mL tube and centrifuged the material for ten min at 3k rpm. We removed supernatant and covered the resulting pellet with 5 mL of RNAlater for storage in the field. In the lab, we extracted samples using the Urine DNA Isolation Kit (Abcam) according to manufacturer protocols. We sheared the resulting elution using the Covaris sonicator as previously described and desiccated the resulting solution down to 20 μL.

### Shotgun Build and Amplification

We built shotgun libraries using the resulting elutions from feces, urine, dentin, and calculus extractions. For initial blunt end repair, we added a total of 20 μL (∼800 ng) of DNA to 5.0 μL NEB Buffer, 0.50 μL dNTP mix (2.5mM), 4.0 μL BSA (10 mg/mL), 5.0 μL ATP (10mM), 2.0 μL T4 PNK, 0.40 μL T4 Polymerase, and 13.10 μL ddH_2_O. We incubated this solution at 15°C for 15 min followed by 25°C for 15 min. We then cleaned the solution using PCR MinElute Purification Kit according to manufacturer protocol before eluting into 18 μL EB buffer. For adapter ligation, we added 18 μL of template DNA to 20 μL Quick Ligase Buffer, 1.0 μL Solexa Mix (Meyer & Kircher, 2010), and 1.0 μL Quick Ligase and incubated the solution at room temperature for 20 min. We then cleaned again using PCR MinElute Purification (Qiagen) according to manufacturer protocol and eluted the solution into 20 μL EB buffer. For the final fill in portion of the shotgun build, we added 20 μL of template DNA to 4.0 μL Thermo pol buffer, 0.50 μL dNTP mix (2.5mM), 2.0 μL Bst polymerase, and 13.50 μL ddH_2_O. We incubated the solution at 37°C for 20 min followed by 80°C for 20 min. We amplified shotgun libraries using Amplitaq Gold before splitting libraries into four identical PCR reactions which contained 9.0 μL of DNA, 9.27 μL PCR Buffer II (10x), 9.27 μL MgCl_2_ (25mM), 3.68 μL dNTP mix (10nM), 2.21 μL BSA (10 mg/mL), 2.0 μL P5 primer, 2.0 μL P7 primer, 61.09 μL of ddH_2_O, and 1.48 μL of Amplitaq Gold enzyme. We used the following PCR conditions: initial denaturation at 95°C for 15 min, followed by cycling of 95°C for 30 sec, 58°C for 30 sec, and 72°C for 45 sec, with a final elongation of 72°C for ten min. We amplified each sample between 8 and 13 cycles (Table S1) using Illumina adapter primers with unique forward and reverse barcodes. We then purified samples using the Minelute PCR Purification Kit according to manufacturer protocol before eluting into 30 μL of EB buffer. We used a total of 7 μL of amplified calculus, dentin, and fecal DNA for each of the capture sets. For urine, we desiccated amplified material from 30 μL down to 7 μL before undergoing a single exome capture.

### Whole-Genome and Exome Capture Kits

We used two whole-genome kits (chimpanzee and human baits) and one human exome kit to capture host DNA from the variety of samples. For the whole-genome chimpanzee kit, Arbor Biosciences produced a custom whole-genome capture MYBaits kit using *Pan troglodytes schweinfurthii* DNA. Genomic DNA extracted from the blood of a chimpanzee (Stone et al., 2010) was used as source material for baits. We pooled extractions for a total of 5 ug of DNA which Arbor Biosciences then used to produce the whole-genome capture baits. For the human whole-genome capture baits, we used a MYcroarray whole-genome human capture kit (using African/Masai male DNA). Finally, we also used the IDT xGen Exome Research Panel (v1.0), a commercially available human exome capture kit.

For feces, we used an input total of 7 uL of amplified material regardless of concentration for each of the three capture kits: the *P. t. schweinfurthii* MYBaits capture, the human MYBaits capture, and the IDT xGen Exome Research Panel. For the chimpanzee whole-genome capture MYBaits kit, we captured each sample according to MYbaits Kapa HiFi HotStart ReadyMix protocol with a hybridization time of 24 hours and a final post-capture PCR amplification of 14 cycles. We purified all samples post-capture through removal of beads, cleanup using the MinElute PCR Purification Kit, and elution into 30 μL. We re-amplified a second time using identical PCR conditions, with the number of cycles dependent upon the outcome of quantification from a Bioanalyzer DNA 1000 chip (Agilent). We purified all samples post-capture using the MinElute PCR Purification Kit according to manufacturer specifications and eluted into 30 μL.

For the MYbaits human whole-genome kit, we captured each of the four amplified fecal samples in the same manner, using the same amount of starting amplified material. However, during the final phase of the MYBaits protocol, all samples were amplified 14 cycles instead of the usual 12 cycles. As such, no samples were re-amplified post-capture after we confirmed high concentrations using a Bioanalyzer DNA 1000 chip.

For the xGen Exome Research Panel, from IDT, the unique P5 and P7 7 nt barcodes used to identify the amplified samples necessitated custom xGen Universal Blocking oligos from IDT. We used a total of 7 ul of amplified material from each sample (greater than the suggested 500 ng input of DNA) for the capture in accordance with manufacturer protocol. The exception to this was for urine, which we desiccated from a starting volume of 30 μL, due to the initial low concentrations. We amplified each capture pool to 12 cycles using KAPA HiFi Hotstart ReadyMix, purified each using Agencourt AMPure beads, and eluted into 22 μL of EB Buffer (Qiagen) as suggested by the protocol. Lastly, we quantified the samples using a Bioanalyzer High Sensitivity DNA chip and amplified each for six more cycles.

Samples were then pooled (see Table 1 for breakdown) before being sent for sequencing at the Yale Center for Genome Analysis. Samples were sequenced on four different Illumina HiSeq2500 Rapid runs (2×100 paired-end) and an Illumina HiSeq2500 standard run (2×150 paired-end).

### Read Processing, Read Mapping, Variant Calling, and Depth of Coverage

Before mapping reads, we examined read quality using FastQC (v0.11.7; http://www.bioinformatics.babraham.ac.uk/projects/fastqc/) and MultiQC (v1.5.dev0; (Ewels, Magnusson, Lundin, & Käller, 2016)), and trimmed adapters and low-quality sequence from reads using BBDuk (v37.90; https://jgi.doe.gov/data-and-tools/bbtools/) with the following parameters: “ktrim=r k=21 mink=11 hdist=2 tbo tpe qtrim=rl trimq=10 minlength=30”. Using default parameters, we mapped reads to the chimpanzee reference genome (panTro4; (Waterson, Lander, Wilson, The Chimpanzee, & Analysis, 2005)) with BWA-MEM (v0.7.17-r1188; (Heng Li, 2013). We then used SAMtools to fix mate pairings, and sort and index BAM files (v1.7; (Heng Li & Durbin, 2009). Because we sequenced some of the samples across multiple lanes (Table S1), we used Sambamba to merge BAM files from these samples using default parameters (v0.6.6; (Tarasov, Vilella, Cuppen, Nijman, & Prins, 2015). Note that we only merged BAM files within individual, biological material, and sequencing library (i.e., samples from the same individual but different source material or capture method were left unmerged and treated separately, as these were different units in our analyses). Finally, we marked duplicates using Picard (v2.18.10; http://broadinstitute.github.io/picard).

We next called variants on each processed BAM file separately using Genome Analysis Toolkit’s (GATK’s) HaplotypeCaller with default parameters (v4.0.8.1; (Van der Auwera et al., 2013)). We then filtered each VCF using BCFTools (v1.6; (H. Li et al., 2009)). We included sites for which mapping quality >= 20, site quality (QUAL) >=30, and genotype quality >= 30.

Because some of the downstream coverage analyses are affected by differing number of raw reads across samples, we downsampled merged BAM files (without duplicate marking) to 40 million reads. To do so, we used SAMtools view (v1.7; (H. Li et al., 2009)) with the flag “-s *downsample_fraction*”, where *downsample_fraction* is equal to 40 million divided by the sample’s total number of raw reads. Note that for analyses requiring downsampling, we only included samples with 40 million or more reads. We next marked duplicates, as above, using Picard (v2.18.10; http://broadinstitute.github.io/picard). We used downsampled BAM files for coverage analyses, but not endogenous content estimates or variant calling.

To calculate depth of coverage from BAM files, we first used SAMTOOLS view (v1.7; (H. Li et al., 2009) with the flags ‘-F 1024 -q 20’ to remove duplicates and only retain reads with the minimum mapping quality of 20. We then used Bedtools GenomeCov (v2.27.1; (Quinlan & Hall, 2010)) with the flag -bg to output a bedgraph file with coverage statistics. Next, again using Bedtools, we intersected bedgraph files with Ensembl coding sequences (CDS) for the panTro4 genome downloaded from the UCSC Table Browser (Karolchik et al., 2004). Finally, using a custom python script, “Compute_histogram_from_bed.py” (see Data Accessibility), we calculated histograms of CDS depth.

### Analysis

We used the SAMtools stats tool to calculate basic metrics related to fraction of reads mapping, duplicates, etc. (v1.7; (H. Li et al., 2009)) across all sample types. We calculated these metrics both with and without duplicates, and for primary and downsampled BAM files. To remove duplicates, we first used SAMtools view with the ‘-F 1024’ flag, before piping output to SAMtools stats. From these metrics, we estimated post-capture endogenous content as the fraction of reads mapping to the reference genome. To test for statistical differences in post-capture endogenous content among sample sources we used an ANCOVA test in R (R Development Core Team, 2014).

Within R, we generated “reverse cumulative” plots (Reed, Meade, & Steinhoff, 1995) of coverage across CDS for feces vs. urine and exome vs. whole-genome for feces (“plot_coverage.R”; see Data Accessibility). These plots display the proportion of total panTro4 CDS (Y-axis) covered by X or more reads (where X is a value on the X-axis).

Using exome data, we examined genotype concordance between paired urine and fecal samples for three individuals (7150, 7507, 7365), and paired calculus and dentin samples for one individual (7057). To estimate concordance, we ensured that variant calls were made at identical sites in the paired samples. We did this by first using BCFtools merge (v1.6; (H. Li et al., 2009)) with the flag “-m all” to merge the paired (urine and feces, or calculus and dentin) exome VCF files for each individual. We then conducted a second round of variant calling using GATK’s HaplotypeCaller (v4.0.8.1; (Van der Auwera et al., 2013)) as described above, with the addition of the flag “-ERC BP_RESOLUTION” and the merged VCF as an interval file via the “-L” flag. These flags force HaplotypeCaller to call genotypes at the same sites—any site called in either the urine or fecal sample (or calculus or dentin sample) from a given individual—in both samples. We then, for each site, compiled genotype, depth, mapping quality, and genotype quality measures from the newly generated VCFs using the custom Python script “Compare_vcfs.py”. From this compiled dataframe, we removed “random” (containing “_random”) and unplaced (containing “chrUn”) scaffolds. We then used the Python script “Process_dropout.py” (see Data Accessibility) to estimate genotype concordance for paired samples at four different minimum depths (4x, 6x, 8x, and 10x). The script finds all sites passing minimum quality thresholds (minimum depth >= value described previously, mapping quality >= 30, and genotype quality >= 30) in both samples, and from those sites counts the number of sites with shared genotypes, genotypes consistent with allelic dropout, and ambiguous genotypes. We considered genotypes consistent with dropout if one of the two samples was heterozygous, while the other was homozygous for one of the alleles in the first sample’s genotype (e.g., “0/1” in urine and “1/1” in feces would be counted as dropout in feces). Genotypes were classified as ambiguous if they were not shared and did not fit a pattern consistent with dropout; for example, if the urine sample had a genotype of “1/1” while the fecal sample had a genotype of “0/2”.

### Data Accessibility

We deposited raw reads in NCBI’s Sequence Read Archive (https://www.ncbi.nlm.nih.gov/sra) under Bioproject PRJNA508503. We implemented the full assembly and analysis pipeline in Snakemake (Köster & Rahmann, 2012), and managed software using Bioconda (Grüning et al., 2018). All code, scripts, and software environments are available on Github (https://github.com/thw17/Gombe_noninvasive_genomic_methods).

## Results

We processed a total of 14 samples from ten different chimpanzees in Gombe National Park, Tanzania from urine (n=7), feces (n=4), dental calculus (n=2), and dentin (n=1) (Table S1). We then captured and sequenced samples using at least one of the following: undirected shotgun amplification (n=2), MYBaits *Pan troglodytes schweinfurthii* capture (Arbor Biosciences; n=4), MYBaits *Homo sapien*s capture (Arbor Biosciences; n=6), xGen (human) Exome Research Panel (IDT; n=26) (Table S1). In total, we analyzed 38 different combinations of individual, source, and sequencing protocol (Table S1).

Concentrations of extracted DNA varied widely across samples (Table S1). Initially, concentrations ranged from 0.11 ng/μL to 65.6 ng/μL with a single urine sample from 7365 too low to be measured. Sequencing success was similarly variable (Table S2). After merging BAM files from the same sample across multiple runs, we generated between 6.9 and 169.5 million reads per sample and while we successfully produced data for the problematic urine sample (individual 7365), it produced the fewest reads (Table S2). We observed high duplication rates likely resulting from PCR amplification during library construction and capture in most, but not all samples (range from 0.05% to 89.4%; Table S2). In general, exome capture had higher duplication rates than whole-genome capture, which, in turn, had higher duplication rates than shotgun sequencing (Table S2; Figure S1). We also observed a linear increase in duplication rate with an increasing number of mapped reads for whole-genome capture, but not exome capture or shotgun sequencing (Figure S1). After filtration and duplicate removal, we were left with between 1.4 and 26.2 million passing reads per sample (Table S2).

Interestingly, we found that samples ranged in the amount of post-capture endogenous DNA (i.e., DNA from the host after sequence capture, as opposed to other sources) from 33.9% to 99.1% (Figure 1; Table S2). We discovered that this effect was driven by the source of the sample (Figure 1; ANCOVA: F(3,18) = 125.493; p < 0.001). Upon further investigation, a post hoc Tukey test revealed that urine (n =7; mean endogenous percentage = 96.4%) had significantly more endogenous DNA than dentin (p = 0.03; n=1; mean=75.9%), feces (p < 0.001; n= 12; mean= 44.9%), and calculus (p < 0.001; n=6; mean=38.3%).

We evaluated capture success using reverse cumulative plots to assess the proportion of CDS in PanTro4 (i.e., the fraction of sequence in PanTro4 annotated as coding sequence) sequenced at different depths. For all samples, we started with a fixed 40 million reads before duplicate removal. We first used fecal samples to compare exome capture with whole-genome capture, and found that exome capture, despite its higher duplication rate, led to broader and deeper coverage across CDS than whole-genome capture (Figure 2). In addition, when comparing urine and fecal samples (exome capture), urine outperformed feces (Figure 3). Across all urine samples, more than 90% of CDS was captured, while only two fecal samples generated data covering more than 50% of CDS (Figure 3). This pattern became even more pronounced as depth increased; for example, at a minimum depth of 8x, more than 75% of CDS was captured in all urine samples, while all fecal samples fell below 10% CDS covered (Figure 3). Finally, when comparing calculus and dentin, we found more than 85% of CDS was captured for the single dentin sample, with 20% of CDS captured at a minimum depth of 8x (Figure 4). However, less than 25% of CDS was captured in both analyzed calculus samples, which decreased to less than 1% at a minimum depth of 8x (Figure 4).

We measured genotype concordance in the three individuals for which we sequenced at least 40 million reads each for paired fecal and urine samples (Table 1; 7150, 7365, 7507) and a single additional individual for paired dentin and calculus (7057). Likely due to the differences in endogenous DNA content and coverage described above, we obtained very few variant sites (i.e., sites with one or both alleles differing from the reference genome) passing quality and depth filters in feces compared to urine (Table 1). For example, at a minimum depth of 10x, we obtained 227, 368, and 2014 sites from the fecal samples from the three individuals, while we obtained 4952, 93,244, and 115,955 sites from the same individuals from urine samples. In total, we were able to compare between 59 and 1,309 sites depending on the individual and depth threshold used (Table 1). Overall, genotypes were overwhelmingly concordant, with less than 11% of sites discordant across all comparisons. Most discordant sites were consistent with a pattern of allelic dropout—that is, one sample was heterozygous, while the other was homozygous for one of the two alleles present in the first sample. Among these dropout sites, at a minimum depth of 10x, feces exhibited higher rates of dropout than urine in two of our three comparisons (fecal dropout = 2-8% of all sites; urine dropout = 0.8-4% of all sites). We also observed “ambiguous” sites—discordant sites inconsistent with the dropout pattern described above—at 1-3% of all sites (Table 1). For calculus and dentin, we compared between 27 and 291 shared sites and observed calculus as having the highest dropout rates of any source of DNA at depth thresholds of 8x and 10x (17.86% and 18.42%, respectively). Although we observed less dropout in dentin, these rates are comparable to our highest observed dropout rates for feces (7.89% dropout at a depth of 10x in dentin, 7.86% dropout in feces at a depth of 10x for individual 7507).

## Discussion

The development of noninvasive genomic methods is critically important for studying wild populations, particularly those that cannot otherwise be legally or ethically sampled. In this study, we evaluated four biological sources of DNA that can be sampled from wild populations of many taxa: feces, urine, dentin, and dental calculus. Feces and urine may be noninvasively sampled from contemporary living populations, while dentin and dental calculus can often be sampled from skeletal collections of wild populations present in collections at museums and field sites. We assessed the quality of these sources in three different ways. First, we determined post-capture endogenous content, the amount of captured DNA is derived from the host. Next, we evaluated the breadth and depth of sequencing coverage across genomic targets. Finally, we measured the concordance of genotypes between pairs of samples captures from different sources from the same individual.

In regard to post-capture endogenous content, of the four sources, we found that urine samples contained the highest proportion of host DNA. While post-capture endogenous content was similar in calculus and feces (ranging from approximately 30-50%), all urine samples contained more than 95% host DNA. Previous studies have demonstrated both that host DNA is present in urine and can be successfully extracted and amplified (Hausknecht et al., 2007; Hayakawa & Takenaka, 1999; Hedmark et al., 2004; Nota & Takenaka, 1999; Valiere & Taberlet, 2000; Waits & Paetkau, 2005); however, our results show for the first time that urine in fact has an high fraction of host DNA compared to other sources of DNA, like feces, that are far more commonly used in genetic studies of wild animals, and thus is well-suited for genomic analysis. While we measured post-capture endogenous content in the same way across the sources of DNA that we tested, we are unable to determine for certain from this study whether the difference in endogenous content directly reflect raw differences in the fraction of host DNA across sources. We cannot easily envision a process that would cause sources of DNA to differ in endogenous content after capture but not before, but future work could aim to measure pre-capture differences to confirm our results.

We found that these differences in endogenous content meaningfully impact downstream sequencing success, as exome capture and sequencing of urine samples led both broader and deeper coverage across the coding sequence of the chimpanzee reference genome than any of the other sources of DNA. With the exception of a single problematic sample, all of the urine samples captured more than 90% of coding sequence at a depth of 4x or greater (after duplicate removal), despite extremely high duplication rates. This means that, without optimization or any other methodological considerations, our urine samples produced sufficient data for most evolutionary and population genetic analyses. In contrast, not a single fecal, calculus, or dentin sample in our study produced enough data for downstream analyses (Figures 3 and 4). Rather than suggest that any of these sources of DNA are more or less useful for genomic analyses, we instead argue that these results indicate that urine might work well “out of the box” similar to other high-quality sources like blood and other tissues, while the other sources that we tested require additional methodological considerations for use, like the many developments for feces (Chiou & Bergey, 2018; Perry et al., 2010; Snyder-Mackler et al., 2016).

Our analyses revealed that genotypes generated from feces and urine from the same individual were broadly concordant, especially when a minimum depth threshold of 10x was used. Urine fared better generally, with fewer sites ambiguously discordant or consistent with dropout. However, we only had paired fecal and urine samples for three individuals, so these results must be taken as preliminary. Regardless of whether genotypes from urine are comparable or better than those of feces, the low rates of allelic dropout underscore the quality of urine as a source of DNA for genomic analyses. In addition, while we are unable to test it at this time, we hypothesize that urine might be less susceptible to problematic contamination than feces. As discussed above, estimates of the proportion of exogenous DNA in urine before capture are unknown; however, it is well known that feces contain overwhelmingly exogenous DNA (Chiou & Bergey, 2018; Perry et al., 2010; Qin et al., 2010). In addition to the microbiota that dominate feces, fecal samples also contain dietary DNA from food items consumed by the host (Bradley et al., 2007; Clayton et al., 2016). In the case of chimpanzees, food items include a wide array of plant and animal items, including nonhuman primate prey (Gilby, 2006; Hobaiter, Samuni, Mullins, Akankwasa, & Zuberbühler, 2017; Mitani, Watts, & Muller, 2002; Pruetz et al.; Uehara, 1997). Because of the extremely high proportion of microbiota in feces, some sort of DNA capture is required to target endogenous DNA (Chiou & Bergey, 2018; Hernandez-Rodriguez et al., 2018; Perry et al., 2010; Snyder-Mackler et al., 2016; White et al., 2019). However, baits can successfully capture sequence across more than 65 million years of divergence (i.e., across the entire primate order) and much of this captured sequence will map to a reference genome equivalently divergent (Webster et al., 2018). This means that the same baits designed to capture host DNA in the feces will also likely successfully capture DNA from primate prey species and that these contaminant sequences will successfully map to the host reference genome, introducing artifacts into genotyping. This possibility needs to be studied further, but if present in our samples, it would artificially increase our observed rates of allelic dropout for urine (calculated as heterozygous sites in feces that are homozygous for one of the alleles in urine). We thus consider our estimates of allelic dropout in urine to be conservative overall.

Our analyses of genotype concordance in dentin and calculus were limited, as we only had a single individual with data from both sources and we recovered very little usable data in the calculus sample. However, in that comparison, we observed a rate of allelic dropout in calculus more than double that of any other tissue. Our estimates for dentin were similar to feces at our most rigorous depth threshold. These results are consistent with previous research showing that yields and quality of host genetic material are lower in calculus compared to dentin (Mann et al., 2018). Yet, calculus has been used to recover full mitochondrial and nuclear genomes from human calculus samples (Ozga et al., 2016; Ziesemer et al., 2018). We therefore suggest that more work is needed to explore and optimize DNA capture from calculus in wild populations.

Taken together, we suggest using urine as a primary source of noninvasive genomic DNA. However, urine is not universally available in sufficient quantities for collection and extraction, and its availability and collectable volume will vary by organism body size, study habitat, and level of habituation. When using other noninvasive biological materials, our results build on previous research (Chiou & Bergey, 2018; Hernandez-Rodriguez et al., 2018; White et al., 2019) showing that targeting a smaller subset of the genome leads to an increase in usable data. In particular, we argue that exome capture is an ideal option, as it targets a small subset of the genome commonly used in evolutionary analyses and there are commercially available human kits that can be used across the entire primate order (Webster et al., 2018). However, like other methods of DNA capture, exome capture requires additional considerations when working with noninvansive samples. First, a multitude of factors impact the quality of host genomic material in a natural environment, including time elapsed since excretion (DeMay et al., 2013), field/laboratory storage conditions (Nsubuga et al., 2004; Panek et al., 2018), and enzymatic activity (Deagle et al., 2006). Second, depending on sample quality, it may be necessary to undergo repeated extractions for the same sample, along with multiple double stranded DNA library builds and multiple indexing amplifications. Third, a single capture of the indexed DNA library may lead to a higher duplication rate, which has been cited in several studies as being a barrier to inexpensive and accurate host genome capture (Bansal & Pinney, 2017; Ebbert et al., 2016; García-García et al., 2016).

Noninvasive samples have been used across a variety of disciplines for addressing many evolutionary and ecological questions (Beja-Pereira, Oliveira, Alves, Schwartz, & Luikart, 2009) including investigations into dietary niches, social structures, and diversity of endangered animals (Carroll et al., 2018). Chimpanzees, currently listed as endangered on the IUCN red list, are considered to be flagship species and indicators of environmental stressors in the surrounding area (Wrangham, 2008). Thus, noninvasive genomic methods are critical for monitoring the health of wild populations as well as aspects of local adaptation and population history important for conservation management. This is especially important for small, isolated populations such as that of Gombe National Park, for which there is an effort to maintain genetic diversity (Pusey, Pintea, Wilson, Kamenya, & Goodall, 2007). The results of our study highlight urine as a promising and untapped source of DNA for this and other genomic work in not only chimpanzees, but wild populations of other protected species as well.

## Acknowledgements

We gratefully acknowledge Nasibu Zuberi and Dismas Mwacha for their assistance in sample collection; the Gombe Stream Research Center and the Jane Goodall Institute; TAWIRI, COSTECH, and Tanzania National Parks for permission to conduct research; the Leakey Foundation, Arizona State University Institute for Human Origins, and the National Institute of General Medical Sciences (NIGMS) of the National Institutes of Health (NIH; grant R35GM124827 to MAW) for generously providing funding; and Research Computing at Arizona State University for providing computing resources.

## Data Accessibility

We deposited raw reads in NCBI’s Sequence Read Archive (https://www.ncbi.nlm.nih.gov/sra) under Bioproject PRJNA508503 (Ozga et al., 2020). We implemented the full assembly and analysis pipeline in Snakemake (Köster & Rahmann, 2012), and managed software using Bioconda (Grüning et al., 2018). All code, scripts, and software environments are available on Github (https://github.com/thw17/Gombe_noninvasive_genomic_methods).

## Author Contributions

ATO, THW, and ACS designed the study. RSN, ICG, AP, YL, and BH provided samples. ATO completed the lab work. THW analyzed the data. MAW and ACS provided laboratory space and funding. ATO and THW wrote the manuscript. All authors revised and approved the manuscript.

---

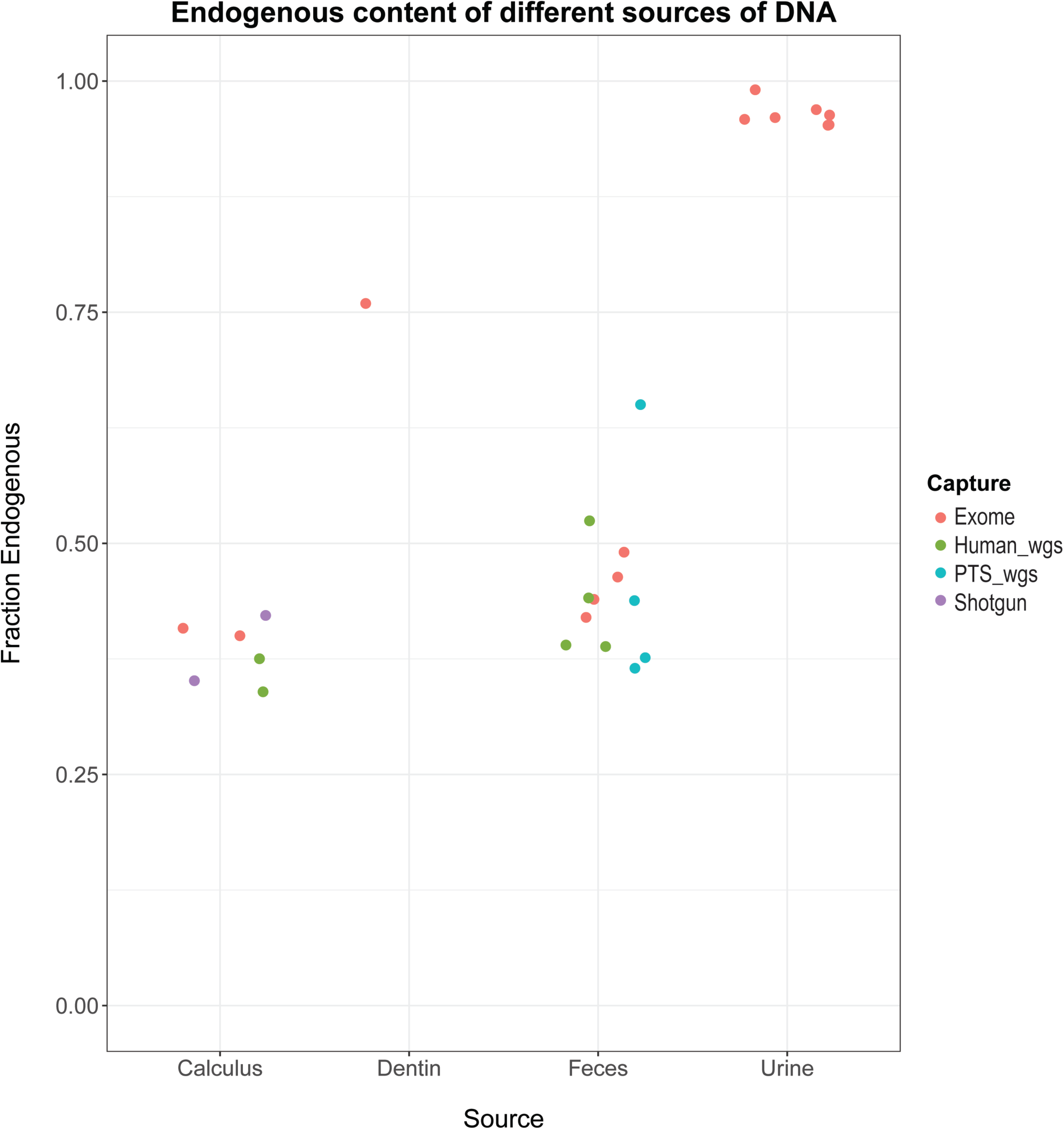

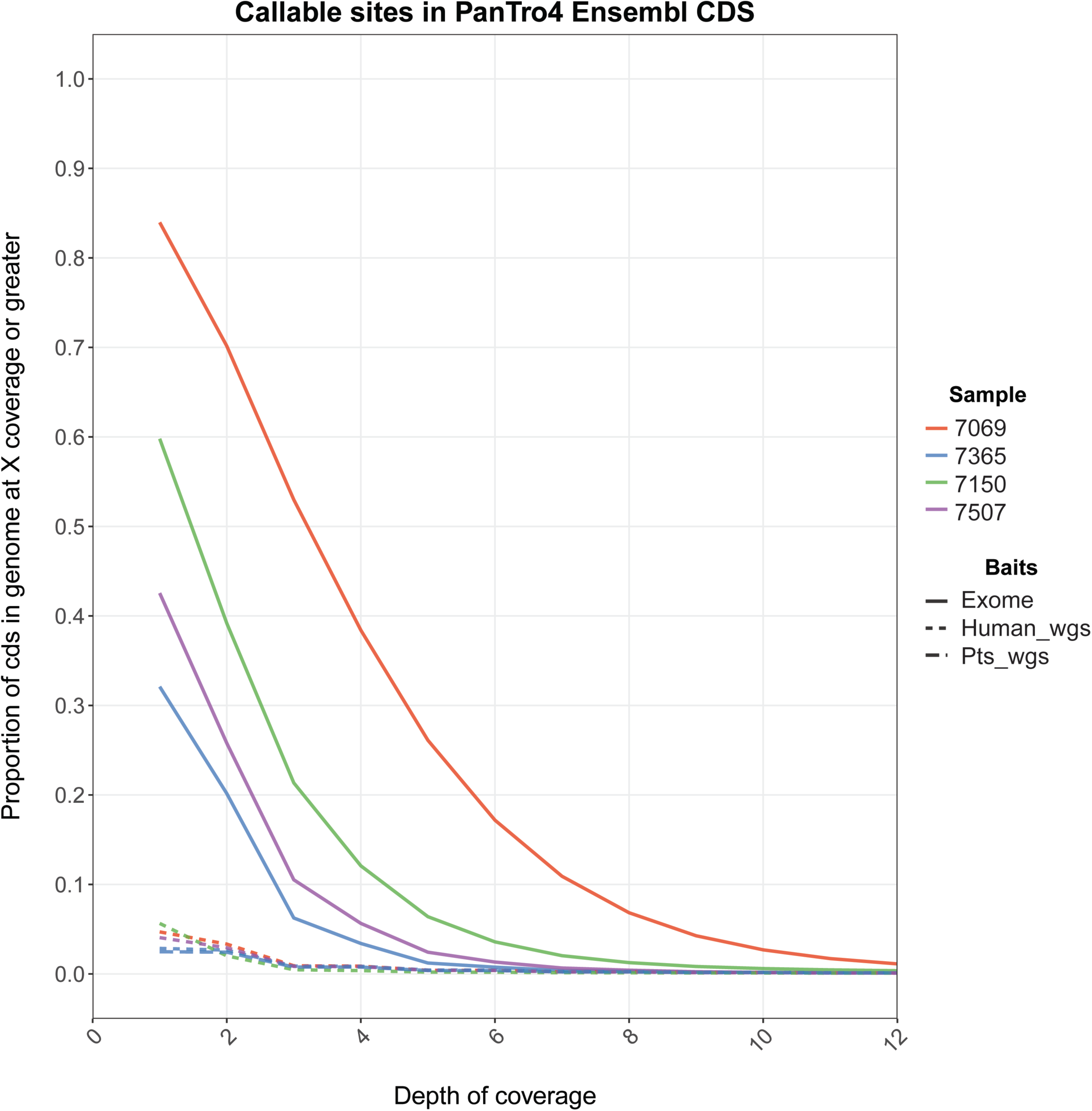

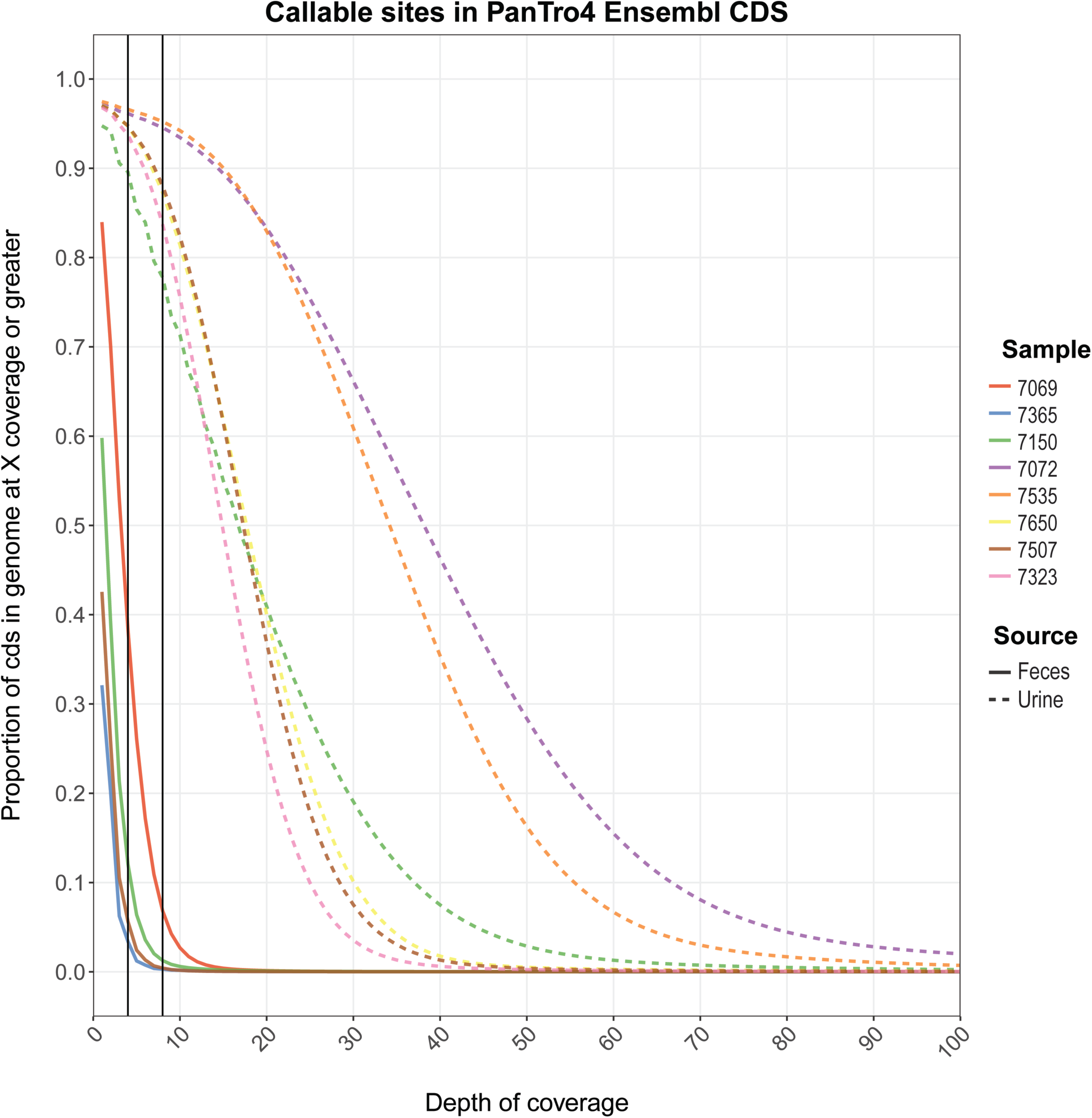

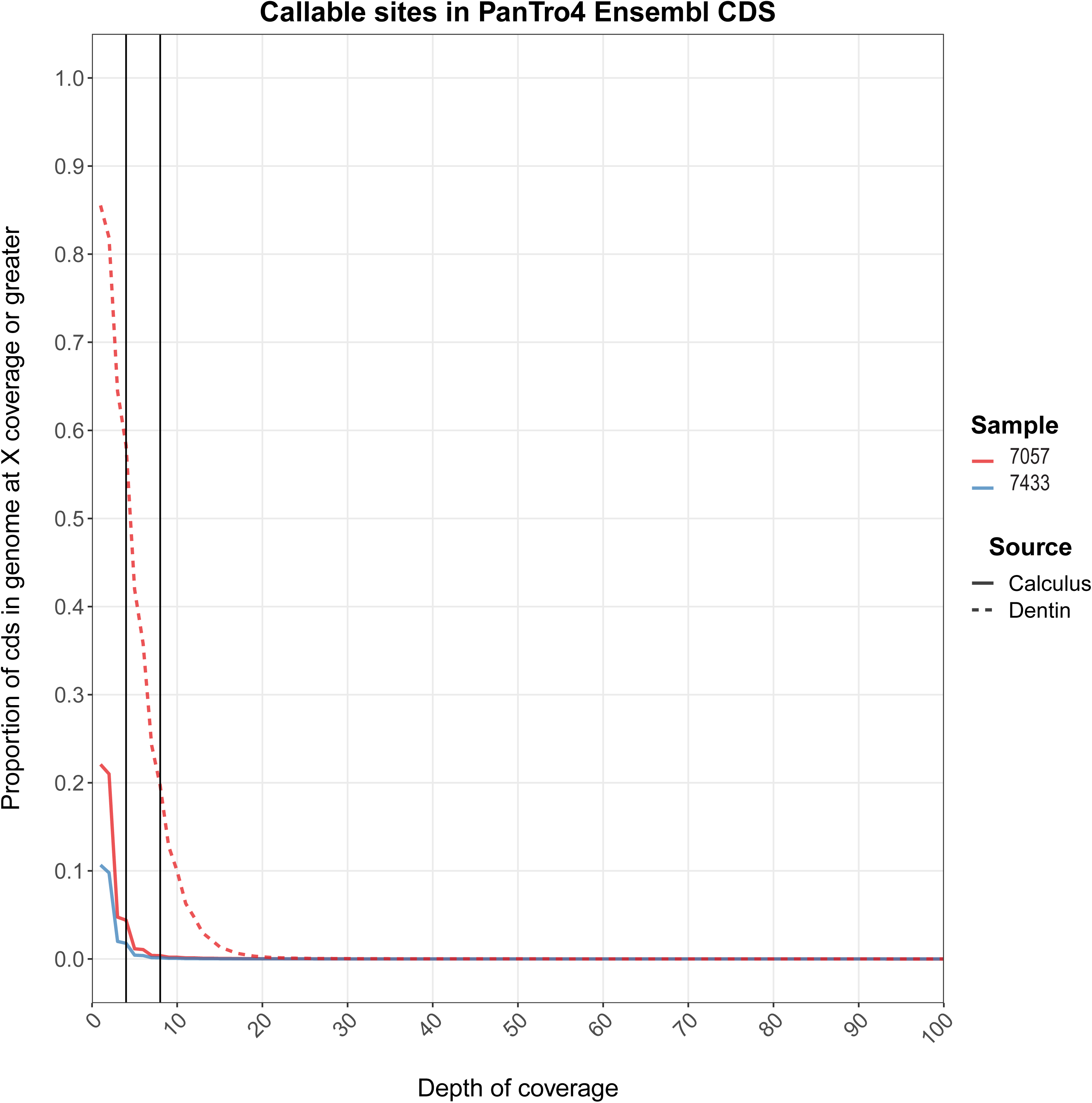

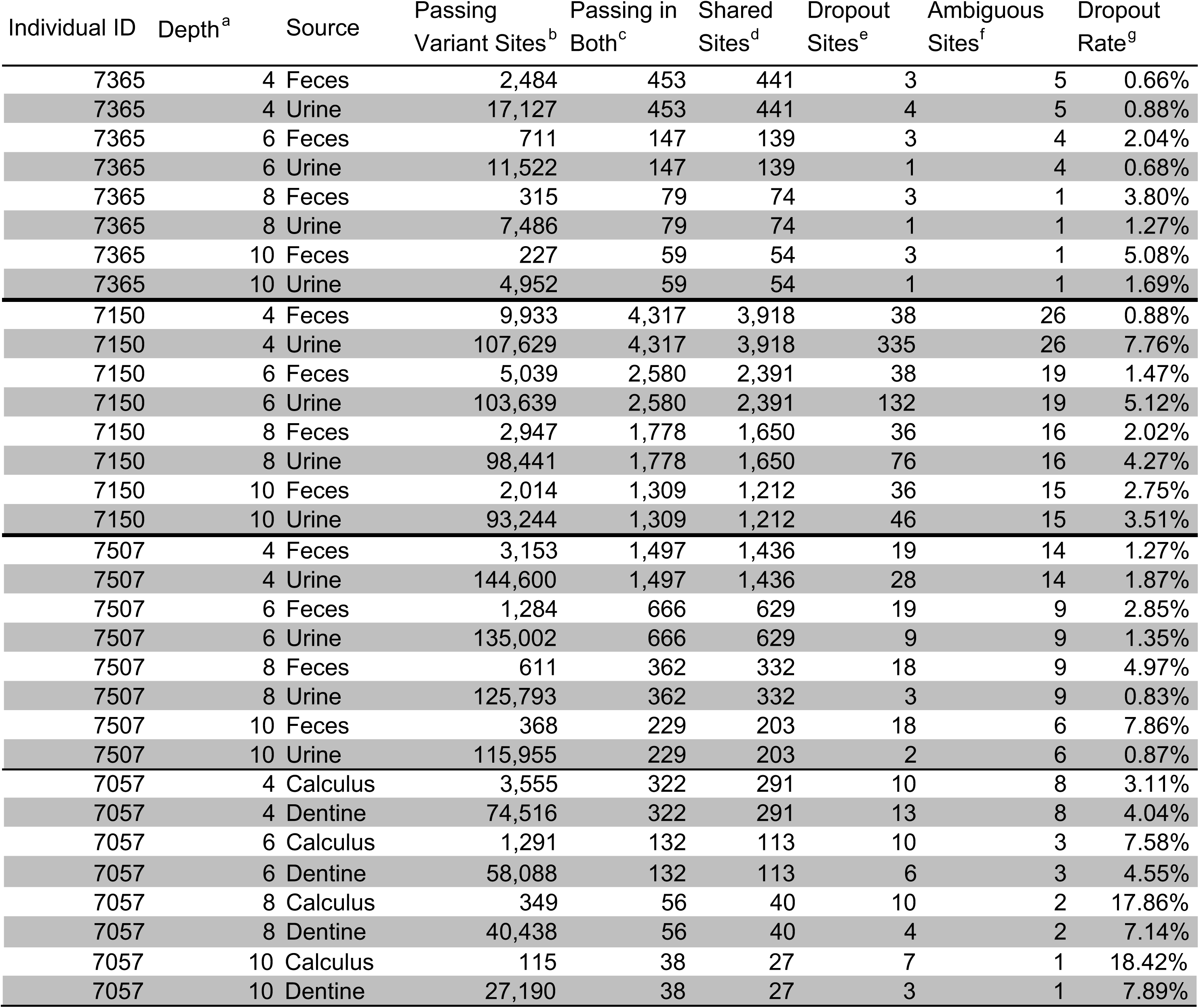

